# A robustly rooted tree of eukaryotes reveals their excavate ancestry

**DOI:** 10.1101/2024.09.04.611237

**Authors:** Kelsey Williamson, Laura Eme, Hector Baños, Charley McCarthy, Edward Susko, Ryoma Kamikawa, Russell J. S. Orr, Sergio A. Muñoz-Gómez, Alastair G. B. Simpson, Andrew J. Roger

## Abstract

The eukaryote Tree of Life (eToL) depicts the relationships among all eukaryotic organisms; its root represents the last eukaryotic common ancestor (LECA) from which all extant complex lifeforms are descended. Locating this root is crucial for reconstructing the features of LECA, both as the endpoint of eukaryogenesis and the start-point for the evolution of the myriad complex traits underpinning the diversification of living eukaryotes. However, the position of the root remains contentious due to pervasive phylogenetic artefacts stemming from inadequate evolutionary models, poor taxon sampling, and limited phylogenetic signal. Here, we estimate the root of the eToL with unprecedented resolution based on a new, much larger, dataset of mitochondrial proteins which includes all known eukaryotic supergroups. Our comprehensive analyses with state-of-the-art phylogenetic models reveal that the eukaryotic root lies between two multi-supergroup assemblages: ‘Opimoda+’ and ‘Diphoda+’. Compellingly, this position is consistently supported across different models and robustness analyses. Notably, groups containing ‘typical excavates’ are placed on both sides of the root, suggesting the complex features of the ‘excavate’ cell architecture trace back to LECA. This study is the most comprehensive phylogenetic investigation of the eukaryote root to date, shedding light on the ancestral cells from which extant eukaryotes arose and providing a crucial framework for investigating the origin and evolution of canonical eukaryotic features.

## INTRODUCTION

Our expanding knowledge of protistan diversity, coupled with the growing availability of genomic data across a broad spectrum of these lineages, has enabled us to reconstruct many high-level relationships within the eukaryote Tree of Life (eToL). Almost all eukaryotic diversity is now categorized into 8-9 prominent ‘supergroups’, with just a small number of known species representing orphan lineages ^1,2^. Despite these advances, resolving the very deepest divergence among living eukaryotes, or ‘root’ of the eToL, remains an ongoing challenge.

The root of the eToL represents the Last Eukaryotic Common Ancestor (LECA), and its correct placement is critical to understanding the nature of LECA, the deepest relationships between supergroups, and the evolution of eukaryote cells, genomes, and life histories. Outgroup rooting is a common method in phylogenetics for determining the root of a tree; by including a distantly related species (the outgroup), the root is identified at the point where the outgroup’s branch connects to the rest of the tree (the ingroup). However, phylogenetic signal for the eukaryote root is difficult to recover, given that LECA existed >1 billion years ago (bya), that large evolutionary distances (‘long branches’) separate eukaryotes from their prokaryotic relatives, and that eukaryote supergroups may have diverged rapidly from one another ^3–5^. Indeed, many previous studies have estimated the position of the root using alternative approaches, such as gene tree parsimony ^6,7^, purportedly primitive molecular and/or cellular features ^8^, gene fusion and domain evolution events ^9,10^, and thought-to-be rare genomic changes ^11^. However, many of these methods lack statistical tests to evaluate their robustness and tend to be sensitive to missing data, incomplete taxonomic sampling, and homoplasy. Importantly, there is also no consensus between them.

Advances in phylogenomic approaches to reconstructing ancient relationships combined with the availability of sequence data from a much more representative set of eukaryotes have led to renewed interest in using outgroup rooting to place the eukaryote root. All extant eukaryotes arose from a chimeric ancestor – some nuclear genes were inherited from a host archaeon related to the Asgard archaea ^12,13^, while other genes are of bacterial origin ^14^; crucially, the latter includes a subset that derives from the endosymbiont related to *Alphaproteobacteria* that ultimately became the mitochondrion ^15,16^. This composite ancestry of eukaryotes makes it feasible to use either a bacterial or an archaeal outgroup to infer the root. Unfortunately, analyses using orthologs of archaeal origin, including a very recent study ^17^, still may be compromised by phylogenetic artefacts that result from a very long branch separating eukaryotes from Archaea. It is well documented that a long branch to the outgroup causes artefactual attraction of long-branched eukaryote lineages to the base of the eukaryote tree ^18^. However, the evolutionary distances (i.e., branch lengths) separating eukaryote genes from their prokaryotic homologs are shorter for genes of ‘mitochondrial’ origin than for those of archaeal origin, potentially making them a more suitable outgroup ^12,14^. Previous attempts at rooting the tree using (alphaproteo)bacterial proteins gave conflicting results; some studies found a root position separating Discoba from all other eukaryotes, while others split eukaryotes into two large clades referred to as Opimoda and Diphoda ^19–22^. However, recent expansions in both the number of curated mitochondrial marker genes ^16^ and relevant novel taxa, along with advancements in phylogenetic modelling, provide unprecedented analytical power to resolve the position of the eukaryote root.

Phylogenetic signal for the position of the root has eroded over time due to multiple amino acid substitutions. Increasing the number of marker genes, and thereby the overall signal, is crucial for a robust rooting inference. A recent study aiming to correctly place mitochondria within *Proteobacteria* identified 108 mitochondrial proteins with alphaproteobacterial affinities ^16^. This is a marked improvement over previous mitochondrial protein datasets used for rooting the eukaryote tree, which had only 39-42 proteins and <12,000 aligned sites ^19,20^. These newly identified marker proteins provide a starting point for developing the largest mitochondrial protein dataset for rooting the eukaryote tree to date. Additionally, high-quality molecular data are now available for many eukaryotic taxa previously excluded from rooting analyses. Some of these, such as Hemimastigophora and Provora, represent newly identified super-group level diversity ^2,23^. Other important taxa absent from prior studies include haptophytes, centrohelids, ancyromonads, telonemids, *Meteora*, and *Anaeramoeba*. Taxonomic sampling across the full breadth of eukaryote diversity is essential both for accurate phylogenetic inference and for drawing appropriate evolutionary conclusions based on the recovered root. For example, if the true root falls between most of eukaryote diversity and a taxon that is missing from the dataset, even a phylogenetically accurate inference of the root would not, in fact, represent LECA.

Evolutionary models commonly used in phylogenetic analyses are often inadequate at capturing the dynamics of sequence evolution on the billion-year timescale. This is known to produce phylogenetic artefacts such as long branch attraction (LBA) in analyses aimed at resolving the most ancient divergences in the Tree of Life ^24^. To address this, many new models of sequence evolution have been developed that account for various evolutionary processes known to confound phylogenetic inference, including site heterogeneity in the substitution process, heterotachy, functional divergence, and compositional heterogeneity amongst taxa ^16,25–29^. These advances in evolutionary modeling are invaluable for rooting the tree of eukaryotes, mitigating potential artefacts and ensuring more accurate and reliable reconstructions of ancient divergences.

Here, we have assembled the largest mitochondrial protein dataset for rooting the eukaryote tree to date, which also includes previously unsampled taxa from across eukaryote diversity. Further, we have applied state-of-the-art evolutionary models for accurate tree estimation and assessed support for alternative root positions while considering potential confounding evolutionary factors. Our analyses robustly support a rooting position deep within the eukaryote tree, placing several supergroups on either side. This study is the most comprehensive investigation of the placement of the eukaryote root to date, offering new insight into the origin of eukaryotes and the ancestor from which they evolved.

## RESULTS AND DISCUSSION

### Curation of a new phylogenomic dataset

To increase phylogenetic signal for estimation of the root, we used the 108 mitochondrial marker proteins identified in Muñoz-Gómez et al.^16^ as a starting point. Proteins with complex evolutionary histories within eukaryotes, such as those affected by numerous duplication and loss events, were excluded, as were those that lacked homologs in most eukaryote groups, resulting in 93 marker proteins being retained (Supplementary Table 1). Eukaryote taxa were selected to represent all major groups, focusing on species with good coverage of the marker protein set. However, including Metamonada was challenging, since this group consists entirely of anaerobes with highly reduced mitochondrion-related organelles (MROs) that lack genomes ^15^. Because of this, metamonads lack most of the genes encoding mitochondrial proteins usually present in other eukaryotes, and most proteins in our dataset could not be found in any available metamonad sequence data. We included two species of *Anaeramoeba* because they were the metamonads possessing the largest number of our 93 mitochondrial proteins. However, *Anaeramoeba flamelloides* and *Anaeramoeba ignava* still possess just 11 and 13 marker genes respectively (16.5% and 19% of the total concatenated alignment sites occupied), and their nuclear genomes exhibit high A+T nucleotide content ^30^. Due to the potential sequence composition biases and lack of signal in the *Anaeramoeba* data, two datasets were generated: one with *Anaerameoba* as representative metamonads (Anae+) and one excluding Metamonada entirely (Anae-). The final datasets retained either 63 (Anae+) or 61 (Anae-) eukaryotes in total. A novel approach based on ancestral sequence reconstruction was used to determine an appropriate size and composition for the outgroup, with 37 taxa being retained (see Methods for details).

Following gene and taxon selection, extensive rounds of manual curation of single gene trees were conducted to exclude any paralogs, contaminants, or horizontal gene transfers. This resulted in a final, concatenated alignment with 98/100 eukaryotic and prokaryotic taxa (Anae-/Anae+), 93 marker proteins, and 22462 sites (Supplementary Table 3).

### Phylogenomic analysis with site-heterogenous models

Both datasets (Anae+ and Anae-) were first analyzed under the site-heterogeneous LG+MEOW80+G4 model; the MEOW80 site profiles were inferred from the Anae+ dataset using a method (see Methods) that extends on the previously published MAMMaL approach ^25^. In brief, this approach separately uses fast- and slow-evolving sites in the alignment to estimate a custom number of site-profile classes (see Supplementary Methods). MEOW80 had the best overall fit for our dataset when compared to other site-heterogeneous models (e.g., LG+C60+G4, LG+UDM64+G4) as assessed by cross-validation (see Supplementary Methods). Although over-parameterization was recently suggested to be a problem for complex site-heterogeneous mixture models ^22^, a thorough investigation found that that tree estimation is generally improved with these site-heterogeneous models and that over-fitting was not a problem, even for very short alignments ^31^.

The maximum likelihood tree under the LG+MEOW80+G4 model recovered most of the generally accepted groups of eukaryotes (Figure 1) with a few notable exceptions including the position of *Mantamonas*, which branches sister to *Thecamonas* (Apusomonadida) rather than within CRuMs ^32^, and *Telonema*, which we do not recover as sister to SAR ^33^. We also do not recover a monophyletic Haptista or Amorphea (Amoebozoa+Obazoa), though paraphyly of the latter is not well supported. Both datasets recover a root separating Amoebozoa, Obazoa, CRuMs, Malawimonadida, Ancyromonadida, and *Anaeramoeba* (for Anae+ dataset only) from all other eukaryotes (including SAR, Archaeplastida, Cryptista, Haptophyta, and Discoba), though support for this root is higher in the Anae− dataset (Figure 1; Figshare). The topologies recovered for the Anae+ and Anae− datasets are identical except that malawimonads branch together with Anaeramoeba as sister to Amoebozoa in the Anae+ dataset, while malawimonads are a sister group to Obazoa in the Anae− dataset, although with low statistical support in both cases. Both datasets were also analyzed with other commonly used mixture models including LG+C60+G4 and LG+UDM64+G4; in all cases, the same root position was recovered (Figshare). The root recovered here is consistent with the Opimoda/Diphoda root proposed by Derelle et al. ^19,20^ but includes many additional major lineages that are newly included here and fall on either side; we henceforth refer to this root position as ‘Opimoda+/Diphoda+’ or ‘OpiDip+’ for short.

**Figure 1.**
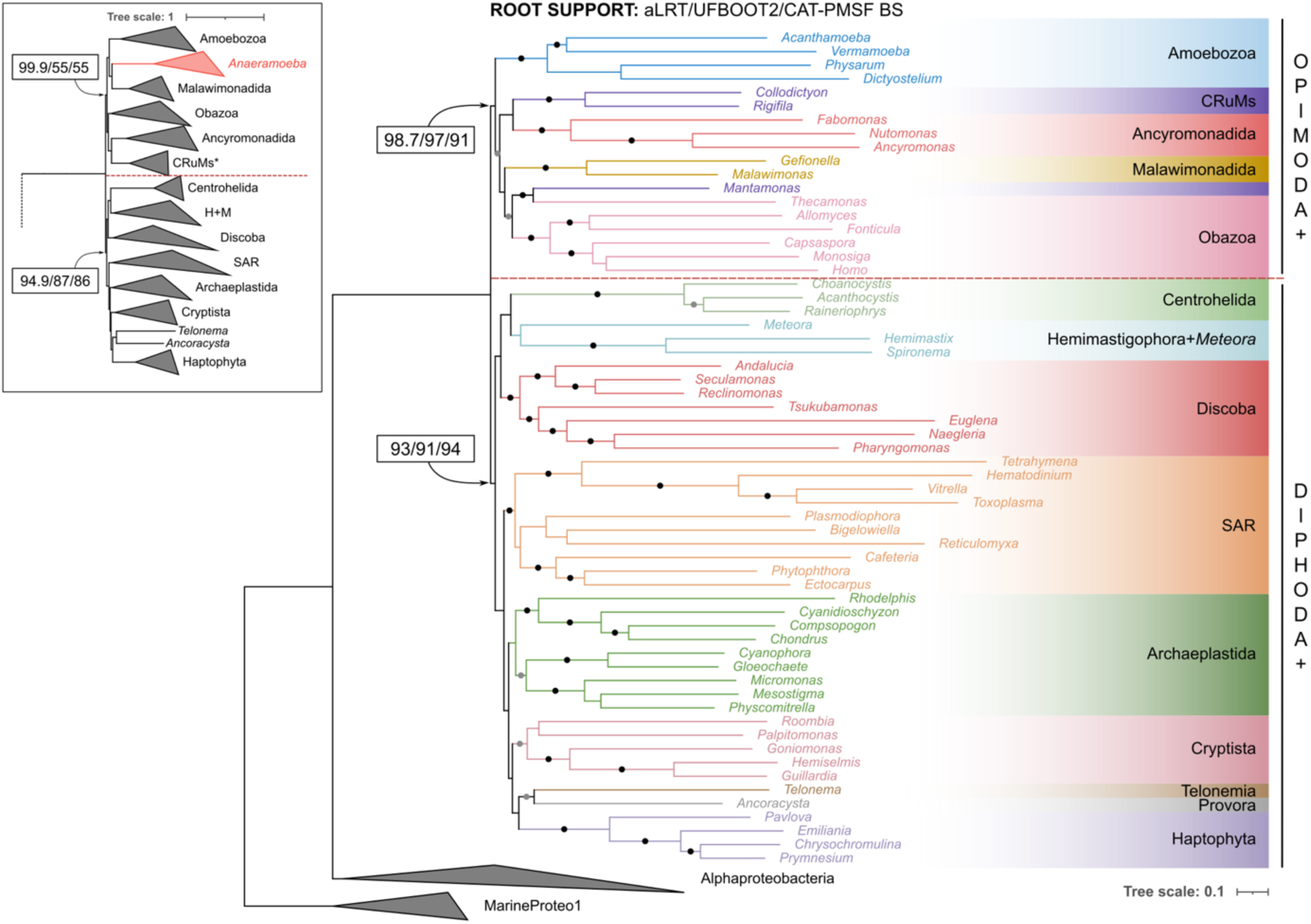
Rooted phylogenies of eukaryotes estimated from mitochondrial proteins of alphaproteobacterial origin. Main tree depicts the phylogeny based on the Anae− dataset, while the inset shows the phylogeny estimated on the Anae+ dataset. Both topologies were estimated with the LG+MEOW80+G4 model. Black and grey circles indicate branches with 100% and ≥85% UFBOOT2 support, respectively. All support values for the two eukaryote root bipartitions are shown, in the order aLRT/UFBOOT2/CAT-PMSF BS. Red, horizontal dashed lines indicate the split implied by the root. Black dashed line in inset indicates branch to outgroup (not shown). Asterisks in the inset signify the placement of *Mantamonas* in Obazoa rather than the expected position within CRuMs.

To evaluate the root in a Bayesian framework, we ran four MCMC chains using Phylobayes under the CAT-GTR model. Unfortunately, the analysis of the Anae+ dataset did not converge, even after 29,000 generations; the consensus tree included the OpiDip+ root, but one chain placed Hemimastigophora on the opposite (Opimoda+) side of the root (Supplementary Fig. 2). Given this failure to converge, we used the recently proposed CAT-PMSF model as an alternative approach ^29^. Here, posterior mean site-specific amino acid frequencies and amino acid exchangeabilities are estimated through Bayesian analysis on a guide tree under the CAT-GTR model and are then used as a user-defined PMSF model in a maximum-likelihood (ML) analysis. Using CAT-PMSF, we also recovered the OpiDip+ root (Supplementary Fig. 3). The full Bayesian and CAT-PMSF analyses were repeated for the Anae− dataset. In this case, the OpiDip+ root was recovered by the posterior consensus tree of all four chains, as well as in the ML analysis with CAT-PMSF (Supplementary Figs. 4 & 5). As with the LG+MEOW80+G4 model, support under CAT-PMSF (bootstrap %) for the root tripartition was higher for the Anae− dataset (91 Opi+/94 Dip+) than for the Anae+ dataset (55 Opi+/86 Dip+).

### Assessing alternative root hypotheses

A variety of eukaryote root hypotheses have been proposed in the literature. These include roots that lie between most eukaryotes and a specific major group, including: i) Archaeplastida ^11^, ii) Euglenozoa ^8^, iii) Malawimonadida ^34^, or iv) Opisthokonta ^6,7^, as well as roots that are basal to (or within) Metamonada or Discoba ^17,21,22,35^. To assess these conflicting hypotheses relative to our optimal inference, we used topology constraints to estimate a set of alternatively rooted trees under the LG+MEOW80+G4 and CAT-PMSF models for both datasets (Figshare). For each analysis, the tested root position was enforced as a topological constraint, with the rest of the topology and parameters optimized by ML.

When comparing the likelihood of the resulting topologies, MEOW80 and CAT-PMSF gave identical rankings of alternative topologies by likelihood. For the Anae+ dataset, our OpiDip+ root had the highest likelihood, followed by roots at the base of *Anaeramoeba* (Metamonada), Discoba, and Malawimonadida, respectively. For the Anae− dataset, the overall order was the same, though an *Anaeramoeba* root was obviously not possible to evaluate (Figure 2; Figshare).

**Figure 2.**
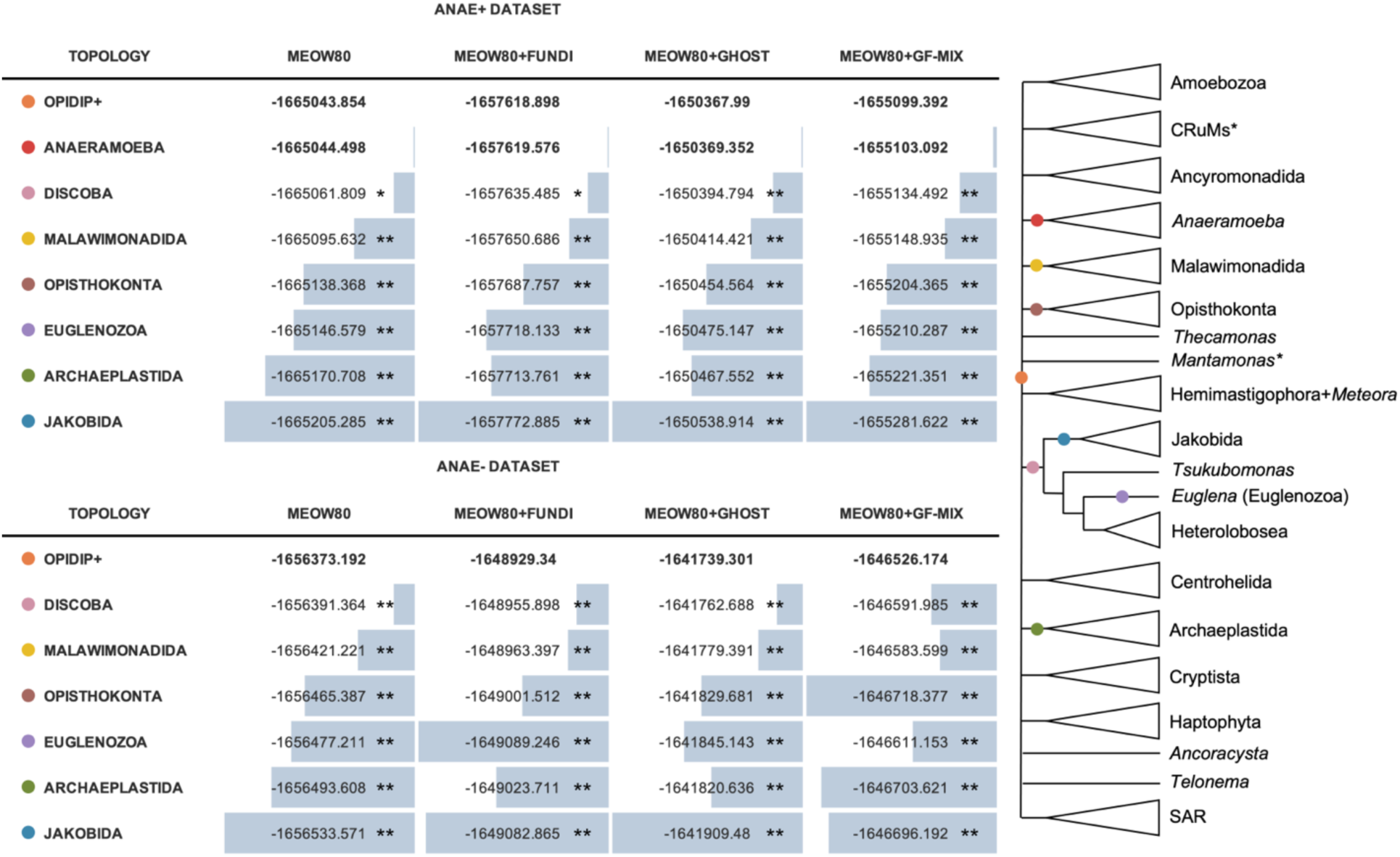
Likelihood estimation of candidate eukaryote roots across complex models. Likelihoods estimated for each fixed candidate topology under all evaluated models for both datasets (with and without *Anaeramoeba*). Likelihoods in bold text are not rejected by either topology test. Those in plain text with a single asterisk are rejected by the Bonferroni-corrected test, while those with two asterisks are rejected by both tests. The multifurcating tree shows the root positions that would be forced under the applied constraints. Each named topology in the table corresponds to a coloured circle in the collapsed summary tree.

### Application of additional complex models

Systematic errors in phylogenetic estimation can result when evolutionary models make incorrect assumptions about the phenomena underlying the substitution process ^24^. Often, this results from assuming homogeneity of truly heterogeneous processes, such as rate heterogeneity, functional divergence, and amino acid compositional heterogeneity. While the models implemented above represent large improvements over earlier site-heterogeneous models, they still do not adequately capture known phenomena involving heterogeneity of the substitution process over branches. To assess whether those factors may be influencing the preferred root, we calculated the likelihoods of the OpiDip+ root and the alternative roots as fixed topologies under three models that account for these phenomena: GHOST ^26^, FunDi ^27^, and GFmix ^16,36^.

The GHOST model captures heterotachy, a phenomenon whereby evolutionary rates vary across sites in a lineage-specific manner, violating the assumptions of the discrete Γ rates model. GHOST is a mixture model comprised of multiple sets of branch length parameters that are optimized by ML ^26^. The FunDi model accounts for sites in proteins that have undergone changes in functional constraints in different lineages ^27^. This process can result in discrete shifts in rates and/or preferred amino acids at many sites on a particular branch of a phylogeny. Finally, the GFmix model was designed to model compositional heterogeneity (changes in amino acid frequencies) both across sites and in different lineages of the tree. For example, it is well-known that many mitochondrial genomes are A+T-biased in their nucleotide composition, leading to increased use of the amino acids F, Y, M, I, N, and K (encoded by A+T-rich codons) and depletion of G, A, R, and P (encoded by G+C rich codons) ^16^. GFmix was recently modified to enable estimation and modeling of composition biases outside of GARP/FYMINK ^36^. In our analyses, no prior assumptions were made as to the composition biases; instead, the groups of amino acids that were enriched versus depleted were estimated directly from the data (see Supplementary Methods).

Using the GHOST, FunDi, and GFmix models, we found that the likelihoods of the various alternative topologies all increase (Figure 2). Both AIC and BIC model selection criteria suggest that the GHOST model provides the best fit to the dataset, followed by FunDi and the base LG+MEOW80+G4 model. AIC and BIC values could not be estimated for GFmix because the branch-specific-composition parameters were not estimated by maximum likelihood. While support for some root hypotheses decreases under some models (e.g., a Euglenozoa root ranks notably worse under FunDi than other models), the OpiDip+ root remains preferred in all cases.

To evaluate the significance of likelihood differences between root positions, we used a likelihood-based *χ*^2^ test with and without a Bonferroni correction for multiple comparisons ^16,37,38^ (Supplementary Table 4). For the Anae− dataset, all alternative topologies other than OpiDip+ are rejected under all models, with and without correction. For the Anae+ dataset, OpiDip+ and *Anaeramoeba* root topologies are not rejected for any model under either test. Additionally, a root at the base of Discoba cannot be rejected using the Bonferroni correction under the base MEOW80 and FunDi models but is rejected in all cases under the GHOST and GFmix models (Figure 2).

Interestingly, for the Anae+ dataset, a root position separating *Anaeramoeba* from all other eukaryotes had nearly the same likelihood as the optimal root under all models and is never rejected by topology testing. When comparing these two root positions as estimated by both MEOW80 and CAT-PMSF, the overall branching orders were identical except for the position of *Anaeramoeba*. However, metamonads are known to represent long branches in phylogenetic trees due to high levels of sequence divergence ^39^, and in our dataset, we recover *Anaeramoeba* as one of the longest branches in the tree. Additionally, as previously mentioned, they also have by far the lowest site occupancy of all taxa in the dataset, providing little signal for their phylogenetic placement. For these reasons, *Anaeramoeba* may be sensitive to the well-known phylogenetic artefact of long-branch attraction (LBA). In LBA, long-branching taxa artefactually branch together when there is a small sample bias ^40^ and/or model misspecification, which could lead to an incorrect placement of *Anaeramoeba* at the long branch connecting eukaryotes to *Alphaproteobacteria*. To test this hypothesis, we used simulation analyses inspired by previous efforts to resolve LBA artefacts in animal phylogenies ^41^.

### Simulation tests indicate an inherent LBA bias towards the *Anaeramoeba* root under realistic model misspecification

A consensus tree was created from the two competing topologies (OpiDip+ vs. *Anaeramoeba*) where any conflicting bifurcations were collapsed to polytomies – this consensus tree was therefore agnostic to either root position. One hundred alignments of the same length as the original alignment were simulated from the fitted CAT-PMSF substitution model applied to the consensus tree with site-specific rates based on estimates from the real data. Gaps were added to the same locations (sites, taxa) as in the original alignment. For each alignment, the likelihoods of the two competing topologies were assessed under three models: CAT-PMSF, LG+MEOW80+G4, and LG+G4. Because the site profiles from CAT-PMSF came from the original data, the simulating model can be considered as close to the ‘true’ model. For data sets simulated from it, the fitted CAT-PMSF model has no model misspecification. The LG+MEOW80+G4 model uses a set of amino acid frequency classes estimated from the original data instead of an independent amino acid frequency profile for each site as in CAT-PMSF and can be considered a case of ‘minor’ model misspecification. On the other hand, LG+G4 does not account for heterogeneity in site-specific amino acid frequencies – this is a ‘major’ model misspecification. If there is no LBA, there should be no preference for one topology over the other (i.e., both topologies are equally likely to be preferred for a given replicate). If there is LBA, there will be a bias toward the *Anaeramoeba* root topology under model misspecification (i.e., the *Anaeramoeba* root topology will have a better likelihood on average).

The difference in log-likelihood between the two topologies (lnL(OpiDip+) – lnL(*Anaeramoeba*)) for each replicate was calculated and overall distributions were compared (Figure 3a). Under CAT-PMSF, we recover an average difference of −0.46 across all replicates. To check whether mean differences from 0 might be due to the necessarily finite number of simulations considered, we conducted two-tailed paired t-tests. For the fitted CAT-PMSF model the average difference was not significantly different from 0 (p-value=0.325). However, under LG+MEOW80+G4, the average difference increases to −4.55, indicating a significant bias towards the *Anaeramoeba* root topology (p-value=0.0002). Under LG+G4, the average value increases to −20.91, indicating greater bias towards an *Anaeramoeba* root (p-value=0.0002). Across replicates, the *Anaeramoeba* root topology is preferred more often than the OpiDip+ root topology with all models, even CAT-PMSF (Figure 3b), and this preference is exaggerated as model misspecification increases. Overall, we see a preference towards an *Anaeramoeba* root topology even with data simulated on a tree that is agnostic to its placement. This indicates that with data like ours, the placement of *Anaeramoeba* is sensitive to LBA artefacts and will preferentially branch at the root of the tree under model misspecification. Repeating this analysis on data simulated in the same way but without missing data results in the same pattern but a greater difference in likelihoods between the two topologies, suggesting this is not due to a small sample bias or any other biases due to missing data alone and that model misspecification is likely playing a role (Supplementary Table 5). Given the potential for LBA bias, poor coverage of *Anaeramoeba* in the alignment, and the lack of signal for its placement in the tree, subsequent analyses focused on the Anae− dataset.

**Figure 3.**
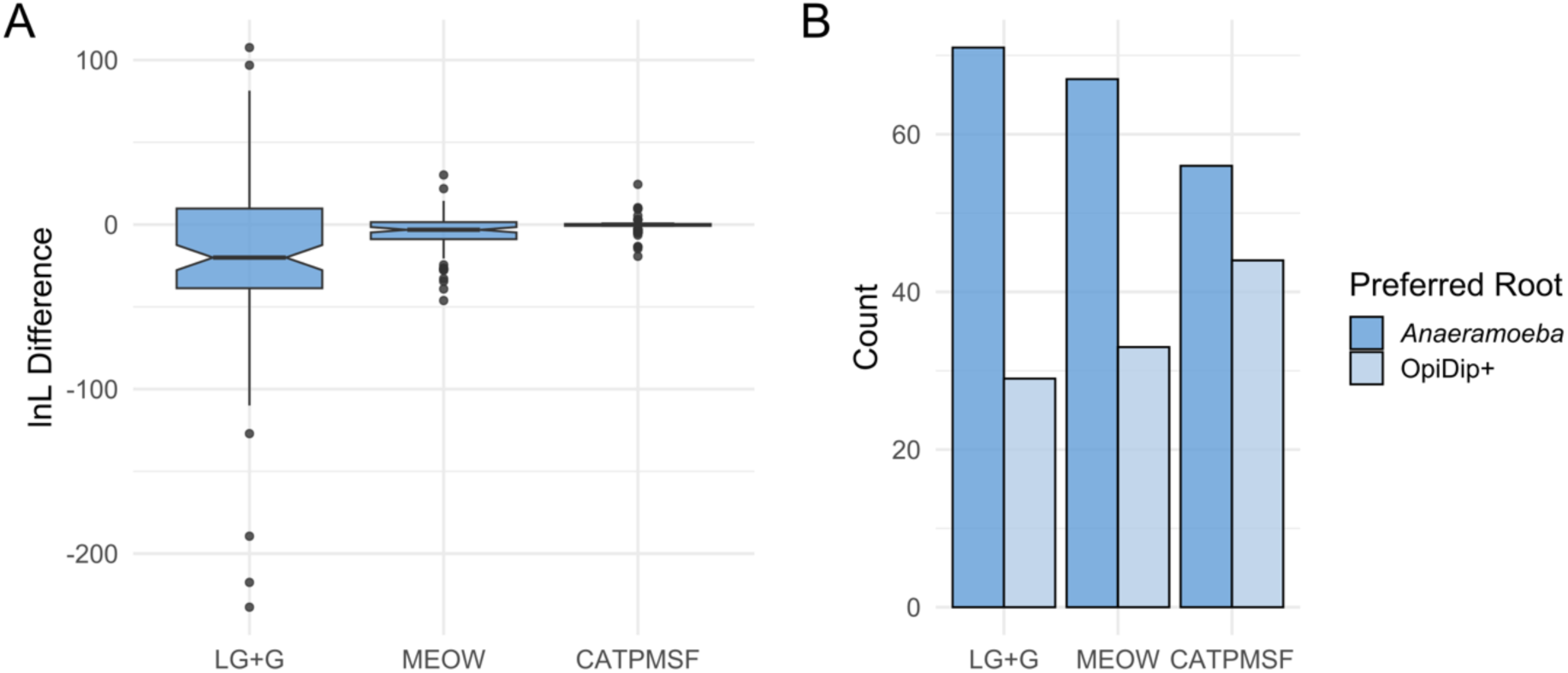
Evaluating potential for LBA bias on simulated data. **A.** Distribution of lnL differences between the two competing topologies (lnL_Opi-Dip+_ - lnL*_Anaeramoeba_*) for 100 simulated alignments across the evaluated evolutionary models. **B.** Total number of simulated alignments that favour either root overall for all evaluated models.

### Removal of divergent sites, genes, and taxa

To further evaluate the robustness of the estimated root and to account for additional potential biases affecting our analysis, several variants of the phylogenomic dataset were generated and assessed. First, the fastest evolving sites were removed in 10% increments until only 50% of the original sites remained. Regardless of the level of site removal, the resulting ML topologies estimated under LG+MEOW80+G4 consistently recovered an OpiDip+ root, albeit with varying degrees of support (Supplementary Fig. 6a). We investigated whether site removal resulted in a stronger preference for another root position by determining the root tripartition recovered in each of the UFBOOT2 replicates for all datasets (Supplementary Fig. 6b). The only other root tripartition to receive >10% support in the UFBOOT2 replicates was a Discoba root, where support ranged from 3.2-37.3%. However, no clear trend was observed in how support varied with increasing site removal.

Additionally, we removed the longest branching taxa from the dataset and analysed the resulting dataset under LG+MEOW80+G4. Again, the root position recovered was consistent with that obtained from the full dataset (Supplementary Fig. 7). Finally, we removed genes exhibiting the longest internal branch; that is, the branch connecting the ingroup (eukaryotes) and the outgroup (*Alphaproteobacteria*). Analysis of the reduced dataset under LG+MEOW80+G4 again supported the same root position (Supplementary Fig. 8). Overall, an OpiDip+ root remains robust when testing for potential biases in sites, genes, and taxa.

### The recovered root suggests LECA was an excavate-like cell

Intriguingly, the topology recovered here places the “excavates” (i.e., Discoba, Metamonada, and Malawimonadida) on both sides of the root. The monophyly of the excavate assemblage is not supported in recent unrooted phylogenomic analyses of eukaryotes, with excavates placed in two or three separate ‘deep’ positions, rather than as one clan ^2,42^. Yet, Discoba, Metamonada, and Malawimonadida each contain ‘typical excavates’ that share a specific complex morphology - a feeding system consisting of a groove supported by a particular cytoskeletal architecture and a vane-bearing posterior flagellum (Figure 4). This shared cell organisation includes some elements also found in certain other taxa distributed across eukaryotes ^43,44^, but also some features without identified equivalents in other groups ^39,45^. If this complex system is homologous in detail across typical excavates, the position of excavate taxa on both sides of the root implies that these traits are ancestral to all eukaryotes; that is, LECA may have been a ‘typical-excavate-like’ cell (Figure 4). Interestingly, the excavate nature of LECA remains very probable regardless of the true phylogenetic placement of *Anaeramoeba* (Metamonada).

**Figure 4.**
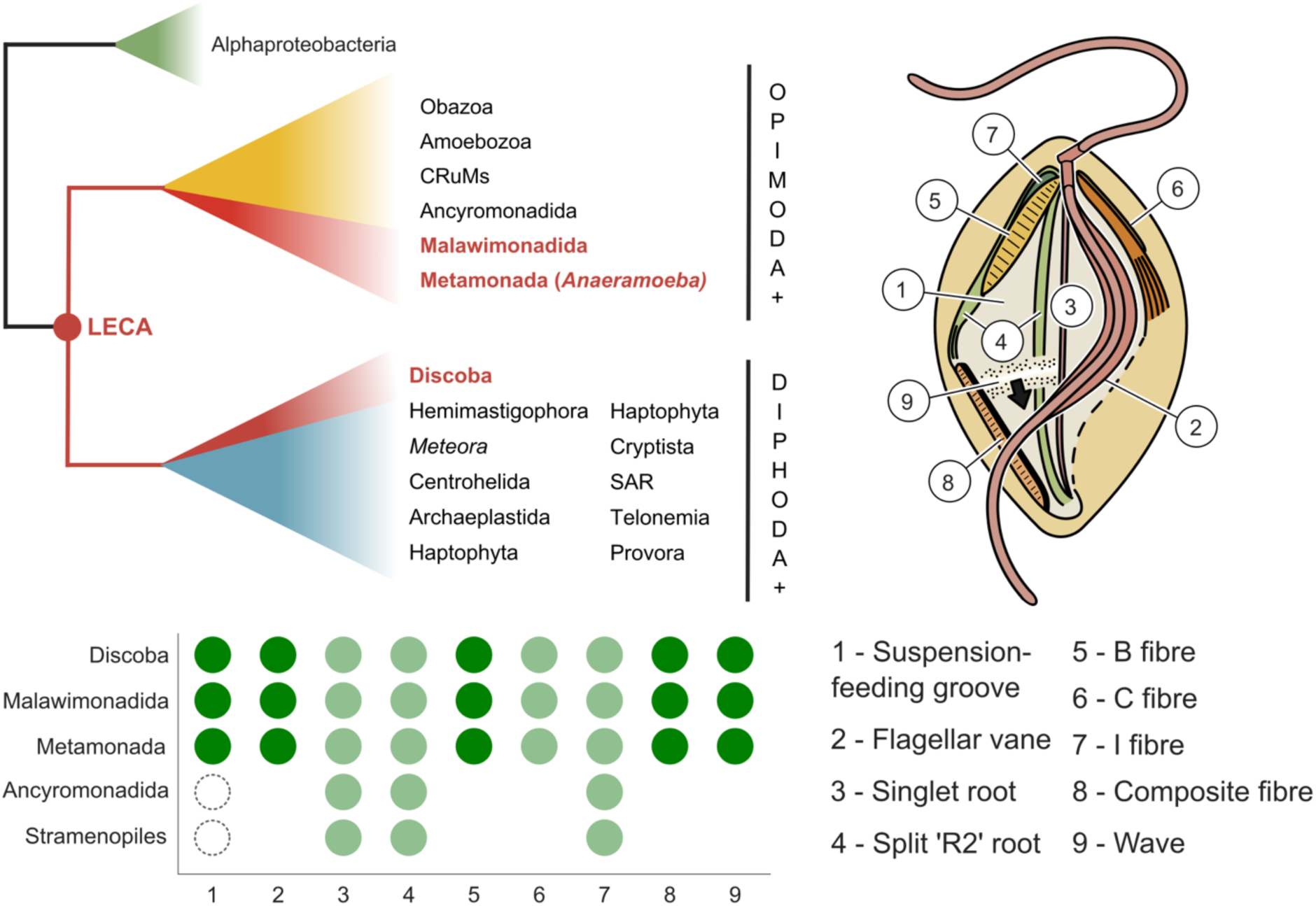
The recovered root suggests the morphological features of ‘typical excavates’ are ancestral traits of LECA. Typical excavates are a non-monophyletic assemblage that shares a particular cell architecture (top right) supported by structures that are either found exclusively in these taxa (bottom left; dark green) or which show a patchy distribution among other eukaryote groups (light green; examples of other groups showing several of these are included). Similar structures found in other groups where homology is unclear are indicated with dashed circles. Flagellar vanes are indicated, provisionally, as specific to typical excavates; somewhat similar vanes are present in selected other taxa, but homology is questionable. The topology recovered in this analysis places the clades containing typical excavates (red) on both sides of the root position, suggesting this architecture to be present in LECA.

This explanation requires multiple losses of the typical excavate cell architecture. However, such losses are known to have occurred inside excavate clades (most clearly, within Metamonada ^46,47^) and the likely deep positions of the excavates within both Opimoda+ and Diphoda+ minimize the number of losses that would need to be inferred. It is also broadly consistent with some of the cell characteristics proposed to unite excavates showing a patchy distribution in other eukaryote groups. For example, parts of the cytoskeletal system supporting the excavate groove appear in deep-branching stramenopiles, ancyromonads, and apusomonads, amongst others ^43,44,48^; in the first two cases, these support a groove-like feeding structure, albeit one that functions very differently than in excavates. Less fully documented grooves are also observed in members of CRuMs ^49,50^, colponemids ^51^, and Provora ^2,52^. While one or more of these grooves may prove to have evolved by convergence, another plausible explanation would be selective retention of the ancestral state during eukaryote diversification.

More generally, the OpiDip+ root supports the view that LECA was a particular kind of complex eukaryotic cell ^53,54^: in this scenario, LECA was likely a relatively small (25 µm or less), unicellular, and flagellated organism. It was phagotrophic and likely bacterivorous, probably feeding on prey in suspension ^45^ rather than as a surface feeder like many amoebae. It possessed mitochondria capable of oxidative phosphorylation ^15^ and had a defined cell shape crucial to its motility and feeding. This shape was underpinned by a complex cytoskeletal organisation, including microtubular roots of diverse sizes, functions, and associated non-microtubular elements.

Models of eukaryogenesis need to have such a form of eukaryote as their “endpoint”. In addition, if LECA was an excavate, it is possible that a suite of cytoskeletal proteins, unidentified at present and potentially absent from animals, plants, fungi, and many other model systems, played important roles in the early diversification of eukaryotes. Many of these proteins could have found new roles in descendant taxa; consequently, even if most eukaryotes are no longer ‘excavates’, their proteomes may carry a legacy of this early phase of eukaryote evolution. Ultimately, the recognition of LECA as a heterotrophic, flagellated excavate will be crucial in studying the processes underpinning eukaryogenesis, and constitutes a major advance in our understanding of the early evolution of complex cells.

## METHODS

### Dataset assembly

Eukaryote taxa were selected for initial inclusion based on the availability of their nuclear (genomic, transcriptomic, or EST) and mitochondrial genomes through either personal communication or publicly available databases such as NCBI or Phylofisher ^55^ (Supplementary Table 6). Taxa without mitochondrial data available were included if they represented major groups that lacked a mitochondrial genome entirely (e.g., Metamonada) or were crucial to ensure diverse representation of all major groups. A few chimeric taxa were created when nuclear genome data was available for one taxon and mitochondrial genome data was available for another, given that they were very closely related to each other (almost always within the same genus; see Supplementary Table 6). Nuclear data were pre-cleaned of bacterial contamination using an in-house script that, for each eukaryotic protein, retrieved the taxonomy associated with the 15 best DIAMOND (v0.9.29) ^56^ hits against nr. If there were more than 13 prokaryotic hits with >85% amino acid identity among those 15 best hits, the sequence was considered probable contamination. Sequences with very few hits in nr were retained at this point. Nuclear and mitochondrial predicted protein sets for each taxon were concatenated.

An initial dataset was provided by Dr. Sergio Muñoz-Gómez (from Muñoz-Gómez et al. ^16^) which contained 111 phylogenetic marker genes from 161 taxa (23 eukaryotes, 138 bacteria). Identification of marker genes is detailed in Muñoz-Gómez et al. ^16^. Hidden Markov Model (HMM) profiles were generated from existing eukaryotic sequences for each marker gene with HMMER (v3.3). If there were less than ten eukaryote sequences for a given marker gene, alphaproteobacterial sequences were added up to ten total sequences before generating the profile. HMMs were used to identify marker gene homologs for each taxon with an e-value cut-off of 1 x 10^-2^ and the top three hits for each were retained. Bacterial and archaeal homologs were added using DIAMOND (v0.9.36) ^56^ with an e-value cut-off of 1 x 10^-3^ against a custom database of 19000 bacterial and archaeal predicted proteomes. All eukaryotic sequences were used as queries and the top 20 hits for each sequence were retained. CD-HIT (v4.7) ^57^ was used with a similarity threshold of 0.8 on the retained prokaryotic sequences to decrease redundancy and overall dataset size.

Alignments were built with MAFFT (v7.471) ^58^ on automatic settings and trimmed with trimAl (v1.4) ^59^ using the -gappyout option. Sequences in the trimmed alignment composed of >70% gaps were removed with BMGE (v1.12) ^60^ unless the total alignment length was >150% the length of known eukaryote orthologs under these trimming conditions (i.e., orthologs curated from the original dataset). This was done to avoid removing reasonably complete orthologs due to much longer homologs (e.g., prokaryotic homologs) being present in the alignment. Single protein trees were estimated with IQ-TREE2 (v2.0.3) ^61^ under the LG+F+G4 model under two conditions: using a full tree search with 1000 replicates of UFBOOT2 ^62^ and using a fast tree search (-fast option) with 100 replicates of true bootstrapping. True bootstrapping support values were mapped on to the optimal tree generated with the full tree search. Single gene trees were manually inspected to remove paralogs, contaminants, and LGTs. Eukaryote sequences that branched strongly (>95% UFBOOT2, >85% true bootstrapping) with archaeal or cyanobacterial clades were discarded as probable nuclear-lineage and plastid homologs. Eukaryote sequences that branched strongly with other prokaryotes or were separated from them by very short branch lengths were discarded as probable contaminants or LGTs. Eukaryote sequences that branched strongly with other eukaryotes they are not closely related to were discarded as potential LGTs or sequencing contamination. Occasionally, two partial sequences from the same taxon were combined to form a single sequence if they branched together with high support and had a region of highly conserved overlap. Once the selected sequences were removed, this process was repeated 5-6 times for each gene until only orthologs remained. However, all subsequent rounds of single gene tree inspection used BMGE (v1.12) for trimming (options: -m BLOSUM30, -g 0.5, -b 3, -h 0.7). Some genes were discarded from the dataset during this process due to their complex evolutionary history (e.g., duplication events that were difficult to parse) or having very few (<5) representative eukaryote orthologs. Following this, sequences were discarded using a custom script if they were <60 amino acids in length to avoid spurious paralog or contaminant inclusion. Extremely long-branching outliers were also removed using a custom Perl script that calculates the average tip-to-tip distance for each taxon and flags taxa that are extreme outliers from the bulk of the data, as defined using Tukey’s fences method ^63^. Briefly, if *x* and *v* are the values of the 25^th^ and the 75^th^ percentiles of the average tip-to-tip distance distribution respectively, then an extreme long-branch outlier has an average tip-to-tip distance > *w* where *w* = 3(*v* − *x*) + *v*. Following curation, a concatenated supermatrix was generated with a custom Perl script.

### Final taxon selection

The size and composition of the outgroup was determined by analysing how outgroup size affects the posterior probability and entropy of the ancestral sequence at the root of *Alphaproteobacteria* as assessed by maximum likelihood. The alphaproteobacteria included in the initial dataset (from Muñoz-Gómez et al. ^16^) were analysed, as they had already been selected to represent known alphaproteobacterial diversity. To minimize the effects of compositional biases on the root position in the final selection of alphaproteobacteria (due to high A+T content in mitochondria and endosymbionts/parasites), the overall GARP/FYMINK amino acid ratios of the marker proteins were calculated separately for a preliminary set of eukaryotic and alphaproteobacterial sequences. Visual inspection of the GARP/FYMINK ratio distributions was used to select 60 alphaproteobacteria whose GARP/FYMINK ratios fell within those most frequently observed in eukaryotes (Supplementary Table 2). Representative taxa across all major groups from *Alphaproteobacteria* were selected, except those where all representatives showed large compositional biases (e.g., *Pelagibacterales*). An outgroup of three representatives of MarineProteo1 and one of *Magnetococcales* was also included to reliably infer the root node. For all marker proteins, alignments were generated with MAFFT (v7.471; --auto) and trimmed with BMGE (v1.12; -m BLOSUM30, -g 0.5, -b 3, -h 0.7). A concatenate generated with only the selected taxa was then analysed using IQ-TREE (v.1.6.12) ^64^ with the LG+C60+F+G4 model. The resulting tree was used as a fixed topology for Treemmer (v0.3) ^65^ to remove alphaproteobacteria 5 at a time while maintaining taxonomic diversity until 9 taxa remained (5 alphaproteobacteria + 4 outgroup taxa), resulting in 12 total datasets. For each dataset, IQ-TREE (v.1.6.12) was used for ancestral sequence reconstruction based on the alignments and fixed tree topologies (option: -asr). For each site at a given node, IQ-TREE outputs the posterior probability of observing each of the possible amino acids at that position. From this, two metrics were calculated for the predicted ancestral sequence at the node connecting *Alphaproteobacteria* to the outgroup:

i. For each site, the character state with the highest posterior probability in the ancestral sequence was considered the ancestral state. The mean, median, and standard deviation of those posterior probabilities across the entire ancestral sequence were then calculated.
ii. For each site, the entropy of the posterior probability distribution was calculated as:

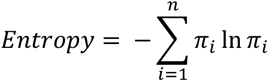

where *π_i_* is the posterior probability of state *i* for all *n* states. The mean, median, and standard deviation of the entropy across the entire ancestral sequence was then calculated.

These values were plotted for all subsets of taxa, and visual inspection of the distribution was used to determine the smallest number of alphaproteobacteria that maintained low entropies and high posterior probabilities of the ancestral sequence to improve computational feasibility (Supplementary Fig. 1a,b).

For eukaryotes, final taxon selection was determined based on the overall gene and site coverage in the final concatenate and to ensure diverse representation (see Supplementary Table 3).

### Site-heterogenous model selection

To estimate a custom profile mixture model from our dataset we used a new approach called “MAMMaL Extension On Whole alignments” (MEOW). This extension of the MAMMaL method ^25^ uses all variable sites in the alignment to estimate a custom number of site-profile classes using user-defined proportions of ‘high-rate’ and ‘low-rate’ sites – see Supplementary Methods for details. Three sets of profile classes were generated using different proportions of high- and low-rate sites – to select amongst these models and other site-profile mixture models, we used a 2-fold corrected Cross-Validation (CV2) procedure ^66^ described in Supplementary Methods. The best fitting model according to CV2 was MEOW(60,20), which will henceforth be referred to as the MEOW80 model.

### Phylogenomic analyses

Posterior mean site-specific frequency distributions of amino acids and posterior mean exchangeabilities for the CAT-PMSF model were generated for the full dataset as described in Szánthó et al. ^29^ using a fixed guide topology estimated under LG+F+G4 in IQ-TREE2 (Figshare). Briefly, two Markov Chain Monte Carlo (MCMC) chains were run under the CAT-GTR model in Phylobayes ^67^ until the effective sample size of all parameters was above 100 before extracting the site-frequency distributions and exchangeabilities for ML analysis.

Maximum-likelihood topologies were estimated in IQ-TREE2 under the following models: LG+MEOW80+G4, LG+C60+G4, LG+UDM64+G4, CAT-PMSF+R4 (Figshare). Mixture weight optimization was enabled (-mwopt). Support was evaluated using 100 replicates of non-parametric bootstrapping under the CAT-PMSF model; trees estimated using all other models were evaluated with 1000 replicates each of UFBOOT2 and aLRT.

Bayesian phylogenies were estimated in Phylobayes using 4 MCMC chains under the CAT-GTR model (Figshare). For the Anae+ dataset, each chain was run for 29,000 iterations with a burn-in of 1000. For the Anae− dataset, each chain was run for 16,500 iterations with a burn-in of 1000.

### Evaluating alternative root positions with additional models

Alternative root topologies were estimated using a constrained tree search in IQ-TREE2. In all cases, three groups were constrained to be monophyletic: the outgroup (*Alphaproteobacteria*) and either side of the tested alternative eukaryote root. All topologies were estimated under the LG+MEOW80+G4 model and all other parameters were optimized freely (Figshare).

The ML topology and all alternative topologies were evaluated as fixed trees in IQ-TREE2 (-te option) with all other free parameters optimized under additional evolutionary models: GHOST and FunDi (Figure 2). For GFmix, the GFmix software was used ^16^. In all cases, the LG exchangeability matrix and MEOW80 site profile mixture models were used as the ‘base’ model. For GFmix and FunDi, the Gamma rate distribution (G4) was used; this was not the case with GHOST, as it cannot be combined with a Gamma rates-across-sites model.

For GHOST, each topology was evaluated with 4, 6, and 8 linked heterotachy classes (LG+MEOW80+H4/6/8) (Figure 2; Supplementary Table 7). For FunDi, functional divergence was assessed across the branch connecting eukaryotes to *Alphaproteobacteria*. Branch lengths and the alpha shape parameter were fixed to the value estimated for the ML tree for each topology. Likelihoods were evaluated for each alternative topology (Figure 2).

We used the modified version of GFmix to estimate the groups of amino acids that tended to be anticorrelated in composition bias across taxa ^36^. This was done separately for different partitions of the data set. To partition the data into compositionally ‘similar’ subsets, compositional heterogeneity within each protein alignment was inferred by first tabulating amino acid counts across alignments for all taxa, including missing data. A chi-squared test was performed on these counts, and Pearson’s residuals were extracted for each amino acid in a taxon. Gaussian clustering of the residuals was then used to partition the dataset into 5 clusters ^68^ (see Supplementary Methods for further details).

Compositional bias was first measured from partitioned sequence data using a binomial test of two proportions ^36^. This requires dividing the taxa into two groups. A chi-squared test was conducted on the amino acid counts across partitions for each taxon and Pearson’s residuals were extracted. UPGMA hierarchical clustering of taxa was performed based on these residuals and taxa were separated into two groups. For each amino acid, a Z-score representing the difference in the count of that amino acid between the two groups was computed using the following equation:

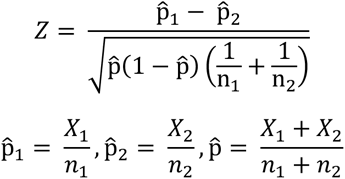

Where X_1_ and X_2_ are the total numbers of that amino acid in groups 1 and 2, and n_1_ and n_2_ are the total numbers of all amino acids in groups 1 and 2. Assuming independence over sites, the Z-score is expected to be distributed like a standard normal random variable under the null hypothesis p1 = p2. |Z| > 1.96 indicates rejection of the null hypothesis at a significance level of p < 0.05. Amino acids with Z > 1.96 were placed into the “enriched” class, amino acids with Z < −1.96 were placed into the “depleted” class. All other amino acids were placed into the “other” class.

To determine the optimal amino acid groups for modeling within each partition, all potential combinations of “enriched” and “depleted” groups of amino acids were evaluated. Chi-squared tests were performed for all enriched/depleted combinations; combinations were ranked by their chi-squared test statistic and the top 50 combinations were retained for GFmix analysis (see Supplementary Methods for more details).

IQTREE2 analysis with LG+MEOW80+G4 was performed for each partition for all topologies, for both datasets. GFmix analysis was performed using the GFmix software (https://www.mathstat.dal.ca/~tsusko/software.html). Hierarchical b-value estimation was performed for each of the 50 enriched/depleted combinations for each tree topology for each partition, and the best-likelihood combination per topology was extracted for each partition. These partition likelihoods were summed together to give full likelihoods for LG+MEOW80+G4+GFmix for each topology (Figure 2).

### Topology testing

Each null hypothesis topology, ***H*_0_**: ***τ*** = ***τ*_0_**, (e.g., an *Anaeramoeba* root is the correct topology) was tested against the alternative topology OpiDip+ 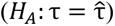 using two different tests. Both tests use the likelihood ratio test statistic (LRS), defined as 2 times the difference in maximized log likelihoods between the two trees: 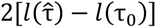. The p-value for the chi-squared test is then calculated as the probability that a chi-squared random variable with d degrees of freedom is larger than LRS, 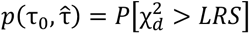, where *d* is the number of 0 edge-lengths in the strict consensus tree of τ_0_ and 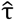.

The chi-squared test was developed as a conservative test for an alternative hypothesis tree that is fixed in advance and independent of the data; it is conservative in the sense that for a 5% test, the type I error rate or probability of false rejection of τ_0_, when true, is less than 5%. When the tree under the alternative hypothesis is estimated from the data, however, that type I error rate can get inflated. To adjust for this inflation, a Bonferroni correction was developed in Muñoz-Gómez et al. ^16^; statistical properties were explored in Markowski and Susko ^38^. As implemented here, the Bonferroni-corrected p-value is calculated as

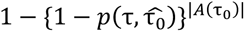

where |*A*(τ_0_)| is the number of topologies that are compatible with the strict consensus tree, excluding those that are also compatible with the root hypothesis; compatible in the sense that the strict consensus tree is a special case with some edge lengths set to 0.

The Bonferroni correction can be very conservative, and the chi-squared test is conservative enough as a test of a fixed-tree alternative that the bias induced via estimation of the tree under the alternative is sometimes not substantial ^38^. Consequently, we report results for both. If the Bonferroni test rejects, then there is very convincing evidence that the tested topology is incorrect.

### Simulation analyses on phylogenetic position of *Anaeramoeba*

A consensus tree was created using the ape library in R from the two competing topologies (OpiDip+ vs. *Anaeramoeba*) where any conflicting bifurcations are collapsed to produce polytomies – this consensus tree was therefore agnostic to either root position (Figshare). Branch lengths for the consensus tree were optimized in IQ-TREE2 under CAT-PMSF+R4. 100 alignments were simulated under the consensus topology using Ali-Sim in IQ-TREE2 (Figshare). Alignments were simulated in such a way that the length of each alignment, the distribution of missing data, the evolutionary rate, and the amino acid frequency profiles for each site were as close as possible to the original alignment. The original alignment was input as a template for alignment length and missing data, while site profiles obtained from CAT-PMSF were supplied to mimic amino acid frequencies for each site. To increase rate variability, the consensus tree was run as a fixed tree under the CAT-PMSF+R14 model, and the most probable rate class for each site was used in simulations.

IQ-TREE2 was run on all simulated alignments on both fixed competing topologies (Opimoda+ and *Anaeramoeba*) given the following models: CAT-PMSF+R14, LG+MEOW80+G4, and LG+G4 (Figure 3; Supplementary Table 8).

### Fast-evolving site, taxa, and gene removal

Fast-evolving sites were identified and removed using a custom Perl script and the per-site rate file output from IQ-TREE2 (.rate file; -wsr option) from the ML topology estimated by LG+MEOW80+G4 (Alignments available on Figshare). Sites were removed in 10% increments and trees were estimated for the resulting alignments under the same model (Figshare).

To approximate the set of taxa that had the longest distance to the root of the tree in the absence of a known root, we calculated the average tip-to-tip distance between a given taxon and the 50% ‘most distant’ other taxa using a custom Perl script and the LG+MEOW80+G4 ML topology. Taxa with an average distance >2.5 were removed from the dataset as potential ‘long branched taxa’ (Supplementary Table 9). The reduced dataset was used to estimate an ML tree under the LG+MEOW80+G4 model in IQ-TREE2 (Figshare).

To determine the genes that have diverged the most between eukaryotes and *Alphaproteobacteria* (i.e., those with the least potential signal for the root), the length of the internal branch connecting eukaryotes and *Alphaproteobacteria* was determined for each gene from single gene trees estimated under the LG+C20+G4 model. These edge-lengths were standardized to have a mean of 0 and variance of 1 over all genes by taking the difference between the branch length and the mean and dividing it by the standard deviation. Standardized edge-lengths larger than 1 were removed (Supplementary Table 10). The trimmed alignments for the remaining genes were concatenated and the ML topology was estimated under LG+MEOW80+G4 in IQ-TREE2.

## DATA AVAILABILITY

All sequence alignments, phylogenetic trees, and custom scripts are available at Figshare.

## ACKNOWLEDGMENTS

We thank N. Ly-Trong and B.Q. Minh for assistance with model implementation in IQ-TREE and J. Jerlström-Hultqvist for assistance with data acquisition and processing. This work was supported by the Moore-Simons Project on the Origin of the Eukaryotic Cell, Simons Foundation grant 735923LPI (DOI: https://doi.org/10.46714/735923LPI) awarded to A.J.R., E.S. and L.E. and by NSERC Discovery Grants awarded to A.J.R. (RGPIN-2022-05430), A.G.B.S. (298366-2019) and E.S. K.W. was supported by a graduate scholarship from the Killam Foundation.

## CONTRIBUTIONS

Ancestral sequence reconstruction analyses, expansion and curation of the final datasets, and all phylogenomic analyses were performed by K.W. in consultation with A.J.R, A.G.B.S., and L.E. K.W. and H.B. performed simulation analyses. H.B. developed and implemented the MEOW model and performed cross-validation testing. C.M. and E.S performed GFmix analyses. E.S. performed topology testing. S.M-G., R.K, and R.J.S.O provided molecular data. K.W, A.J.R, A.G.B.S, H.B., C.M., and E.S. wrote, and all authors edited and approved, the manuscript. A.J.R and L.E initially conceived the study.

