## Supplementary Information for "A robustly rooted tree of eukaryotes reveals their excavate ancestry"

### SUPPLEMENTARY FIGURES

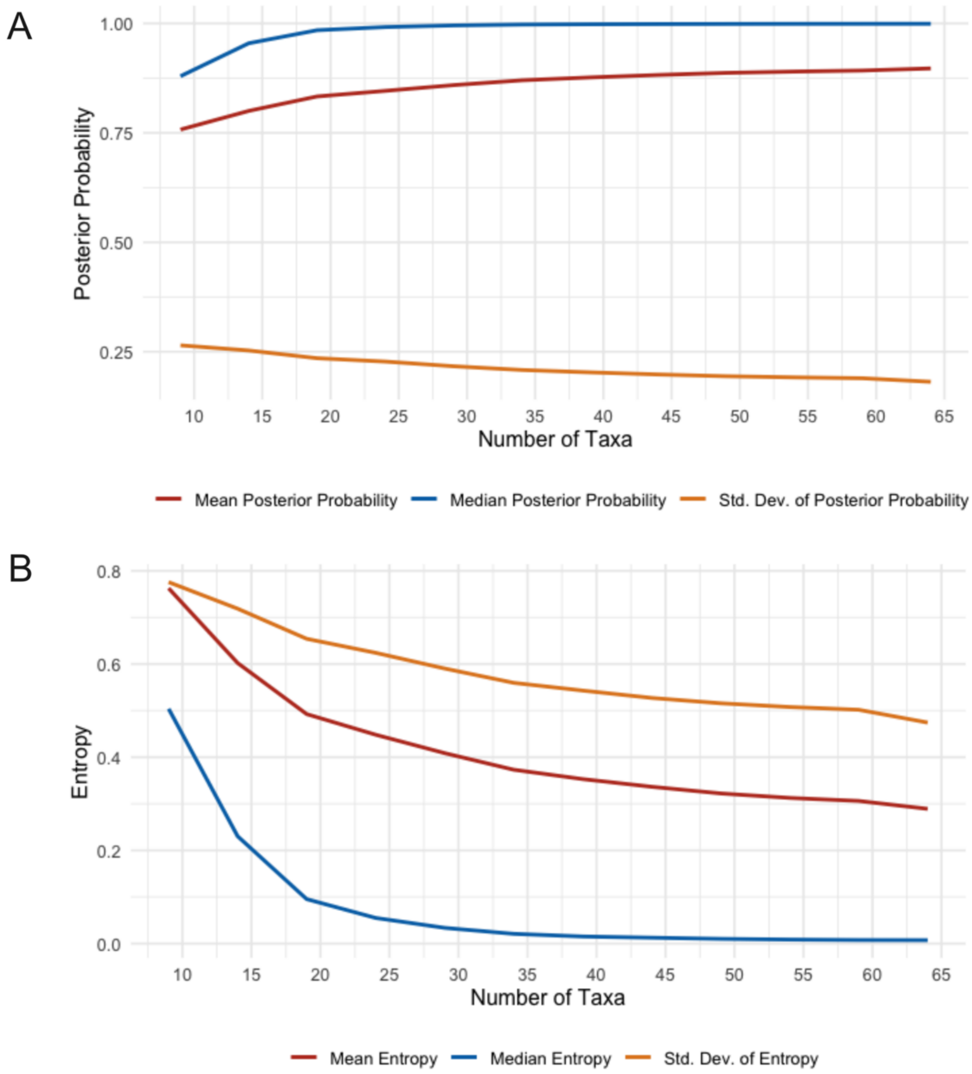


**Supplementary Figure 1. Entropy and posterior probability of the predicted ancestral sequence at the root of *Alphaproteobacteria* as ingroup taxa are removed.** A. The mean, median, and standard deviation of the posterior probability as taxa are removed. B. The mean, median, and standard deviation of the entropy as taxa are removed.

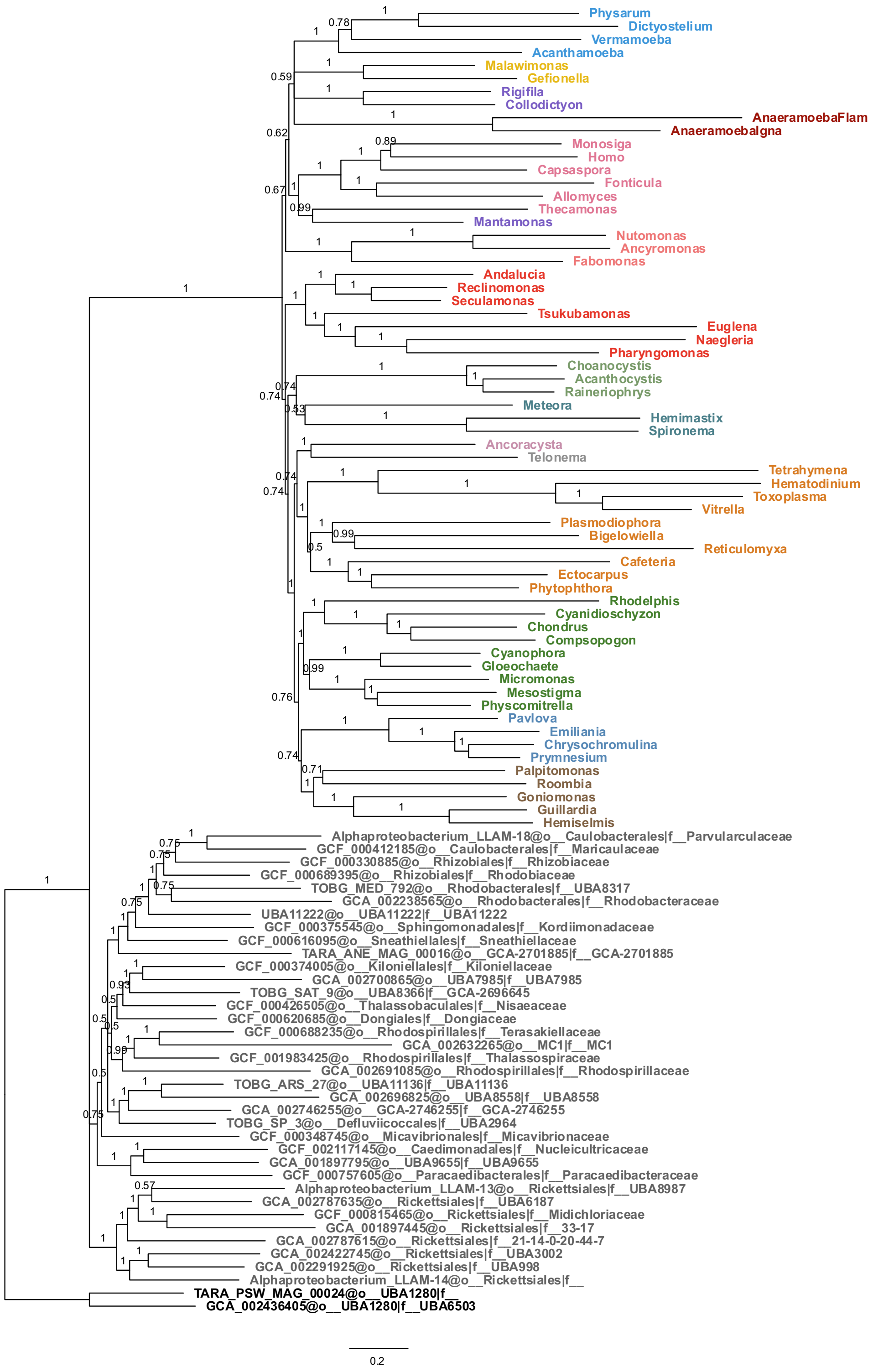


**Supplementary Figure 2. Bayesian consensus tree estimated from Anae+ dataset.** Consensus phylogeny estimated under the CAT+GTR model in PhyloBayes from 4 chains after 29,000 cycles, with a burn-in of 1000. Posterior probabilities for each bipartition are indicated. Note that these 4 chains included one in which Hemimastigophora branched on the opposite side of the root, separate from its previously inferred close relative *Meteora*, with this non-convergence resulting in posterior probabilities ~0.75 (0.74) for several deep nodes in the Diphoda+ side of the tree.

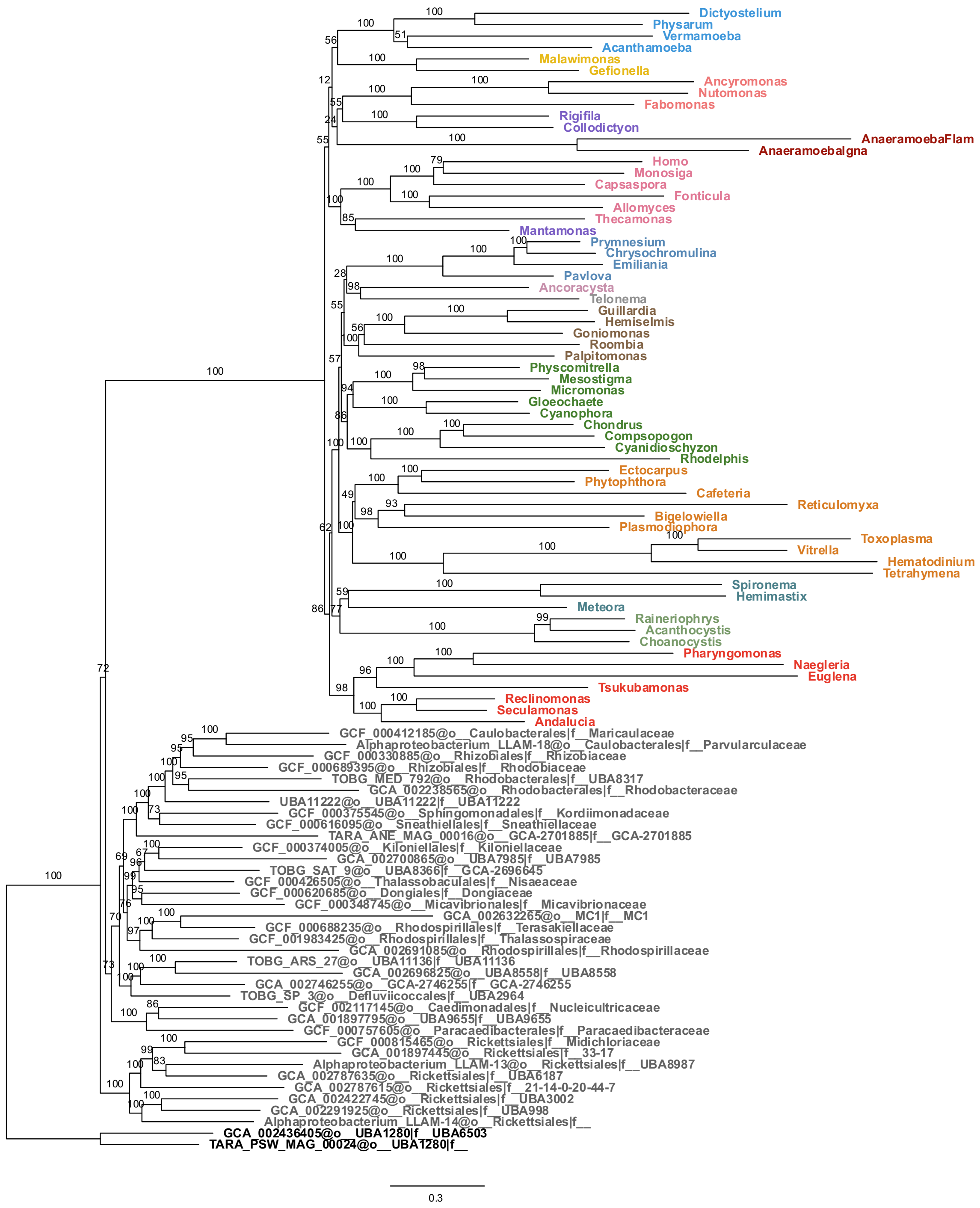


**Supplementary Figure 3. Phylogeny estimated from Anae+ dataset with CATPMSF model.** Amino acid exchangeability matrix and site frequency profiles were estimated on the alignment and a guide tree in PhyloBayes. Rate variation was modelled with the Free Rate model with four classes. Support values indicated are from 100 replicates of non-parametric bootstrapping.

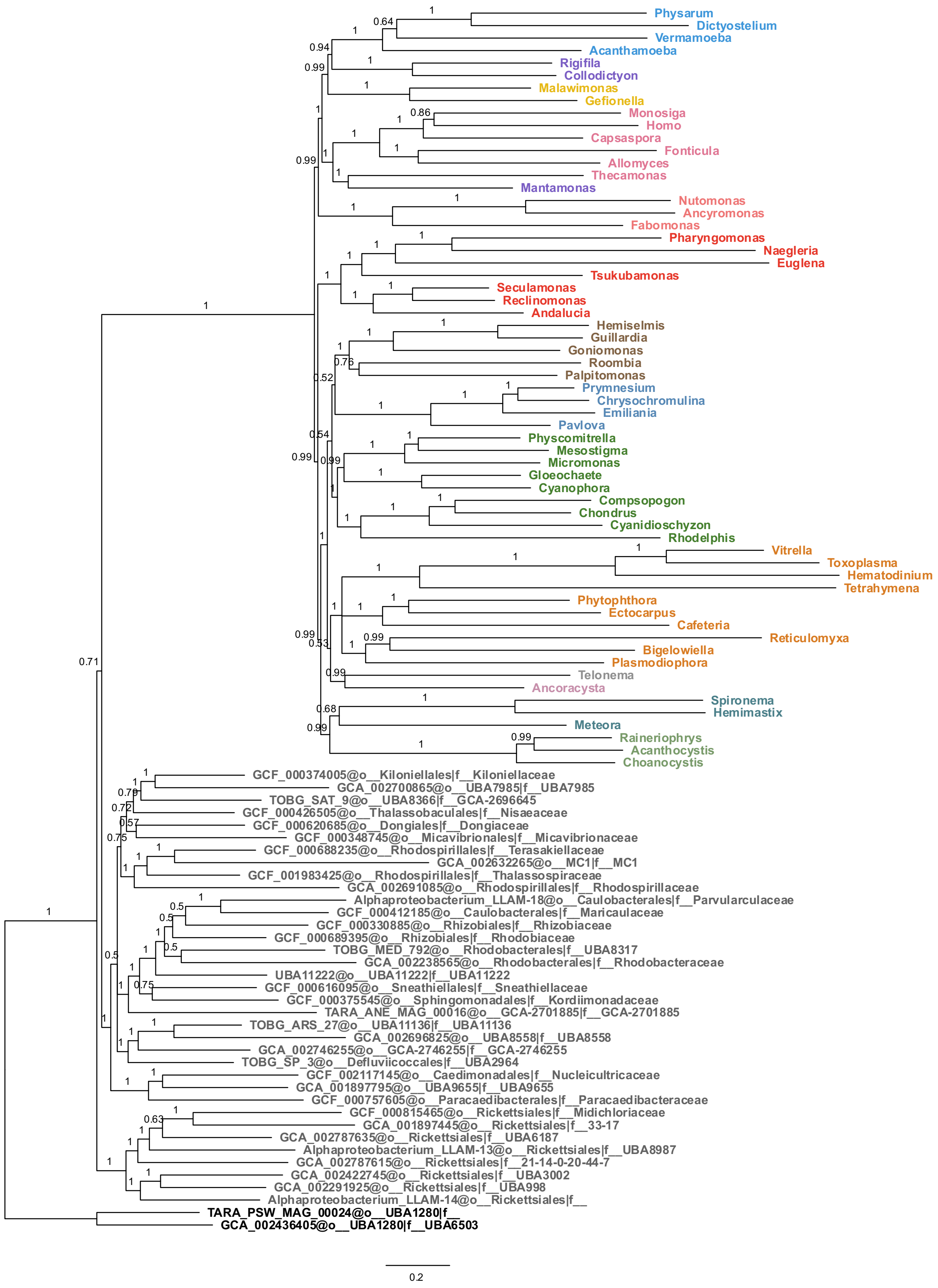


**Supplementary Figure 4. Bayesian consensus tree estimated from Anae- dataset.** Consensus phylogeny estimated under the CAT+GTR model in PhyloBayes from 4 chains after 16,500 cycles, with a burn-in of 1000. Posterior probabilities for each bipartition are indicated.

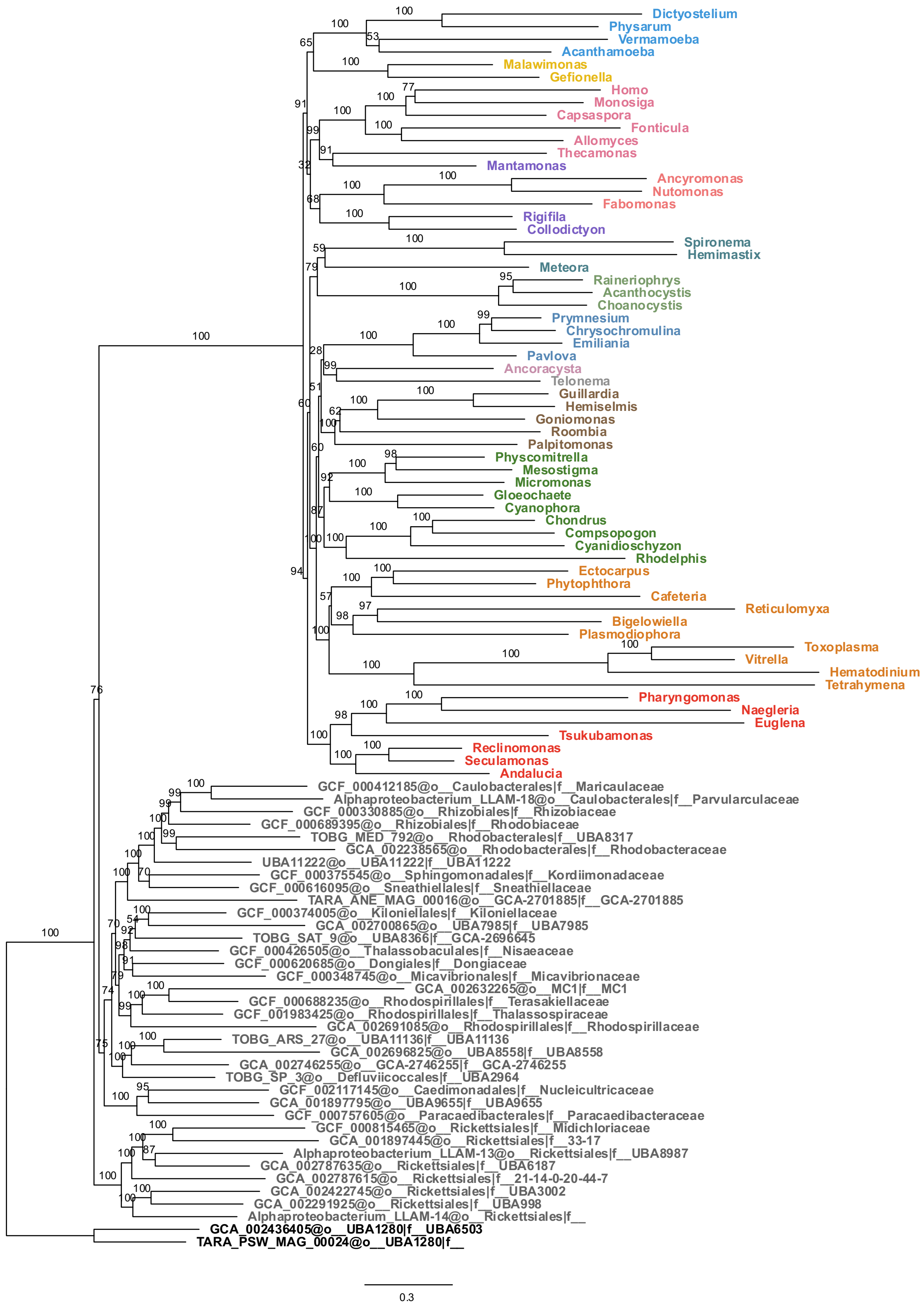


**Supplementary Figure 5. Phylogeny estimated from Anae- dataset with CATPMSF model.** Amino acid exchangeability matrix and site frequency profiles were estimated on the alignment and a guide tree in PhyloBayes. Rate variation was modelled with the Free Rate model with four classes. Support values indicated are from 100 replicates of non-parametric bootstrapping.

A

| % Sites Removed | aLRT/UFBOOT Support | |
|---|---|---|
| | Opimoda+ | Diphoda+ |
| 10 | 98.5/93 | 53.2/75 |
| 20 | 99.4/98 | 78.7/82 |
| 30 | 99.2/93 | 50.3/75 |
| 40 | 94.6/93 | 30.2/55 |
| 50 | 98.0/97 | 95.8/87 |

B

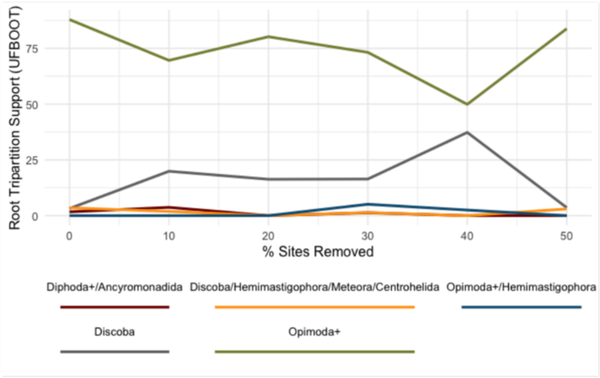


**Supplementary Figure 6. Evaluating change in support for the root position as fastest-evolving sites are removed.** A. Phylogenies were estimated in IQ-TREE2 under the LG+MEOW80+G4 model with 1000 replicates each of aLRT and UFBOOT2. Support values are indicated for the bipartitions relevant to the position of the eukaryote root.

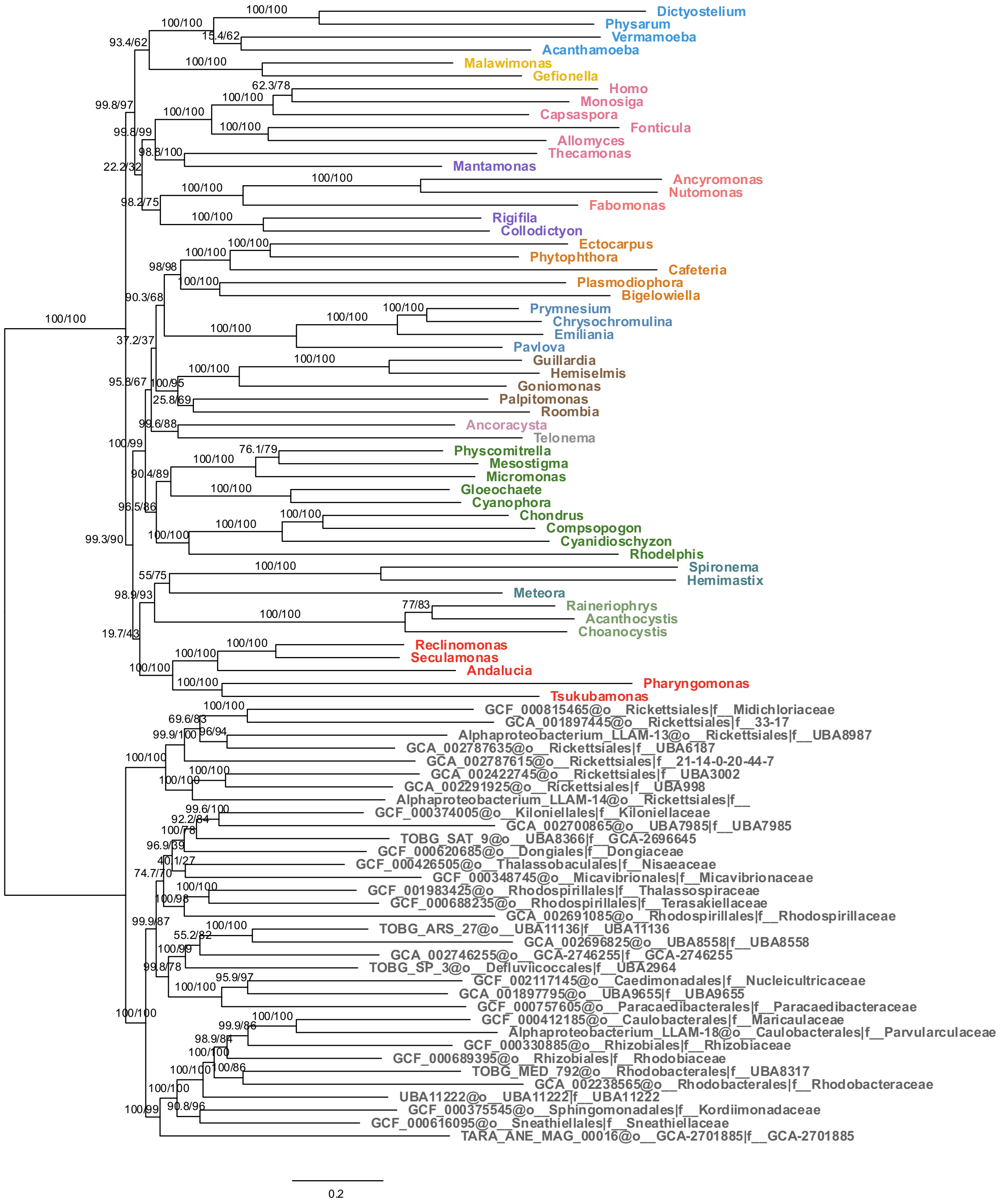


**Supplementary Figure 7. Phylogeny estimated from Anae- dataset after removal of fastest evolving taxa.** Phylogeny was estimated in IQ-TREE2 under the LG+MEOW80+G4 model with 1000 replicates each of aLRT and UFBOOT2. Support values are displayed on branches as aLRT/UFBOOT2.

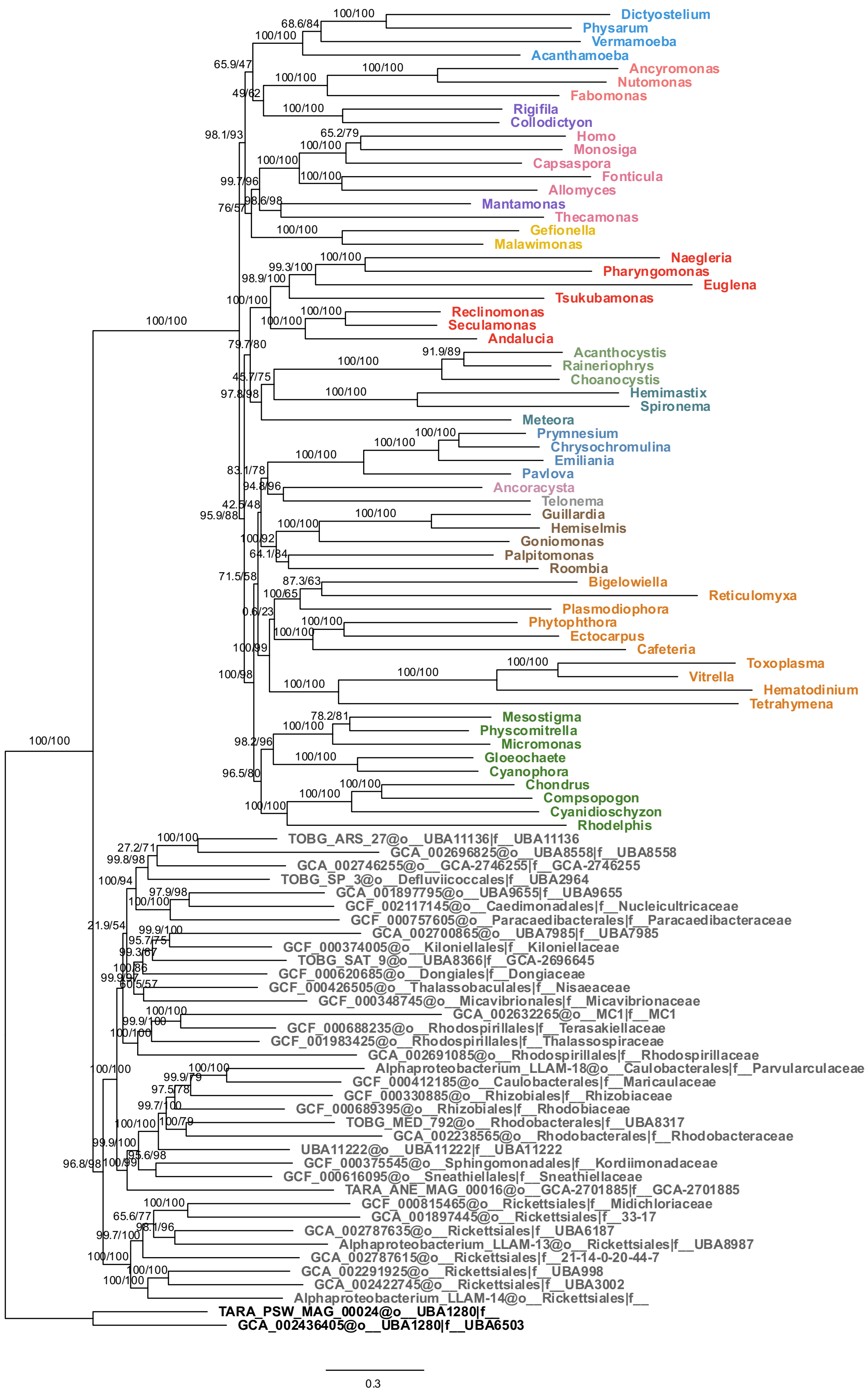


**Supplementary Figure 8. Phylogeny estimated from Anae- dataset after removal of most divergent genes.** Alignment after gene removal consisted of 19324 sites from 80 concatenated genes. The phylogeny was estimated in IQ-TREE2 under the LG+MEOW80+G4 model with 1000 replicates each of aLRT and UFBOOT2. Support values are displayed on branches as aLRT/UFBOOT2.

### SUPPLEMENTARY METHODS

**Custom profile mixture model estimation with MEOW.** The MEOW model is an extension of the MAMMaL method [1] that uses all variable sites in the alignment to estimate the site-profile classes. First, invariant sites in the alignment were removed from consideration and the site rates of the variable sites were estimated based on a guide tree (estimated using the LG+C60+G4 model with IQ-TREE) using the Discrete Gamma Probability Estimate (DGPE) method of site rate estimation [2]. Sites were then partitioned into a set corresponding to the top 25% of rates ("high-rate sites") and the bottom 75% ("low-rate sites"). Site profile classes were then estimated separately from the high-rate sites and the low-rate sites using the MAMMaL approach. The final set of 80 site profile classes were generated by combining site profile classes estimated from these two sets of sites. Three different final sets of profile classes were generated by estimating different numbers of classes from each site-rate set: MEOW(80,0) estimated 80 classes from high-rate sites only, MEOW(40,40) estimated 40 classes from each of the high- and low-rate site sets, and MEOW(60,20) estimated 60 classes from high-rate sites and 20 from low-rate sites.

**2-fold corrected cross-validation.** To select the optimal profile mixture model for our data we used the 2-fold corrected Cross-validation procedure (CV2) referred to as the Burman correction in Susko & Roger [3]. For an alignment $\boldsymbol{X}$ divided into two equal-sized non-overlapping sets of sites $\boldsymbol{y}$ and $\boldsymbol{z}$, the 2-fold cross-validation score is given by

$$lnL\left(\boldsymbol{X}, T, \hat{\theta}(\boldsymbol{X})\right) - \hat{\beta}$$

Where $lnL(X,T,\hat{\theta}(X))$ is the maximized log-likelihood for the whole alignment $\boldsymbol{X}$ and tree $T$ and adjustable parameters $\theta$ and

$$\hat{\beta} = \frac{1}{2}\left\{lnL[\boldsymbol{y}, \hat{\theta}(\boldsymbol{y})] - lnL[\boldsymbol{z}, \hat{\theta}(\boldsymbol{y})] + lnL[\boldsymbol{z}, \hat{\theta}(\boldsymbol{z})] - lnL[\boldsymbol{y}, \hat{\theta}(\boldsymbol{z})]\right\}$$

Where $lnL[\boldsymbol{y}, \hat{\theta}(\boldsymbol{y})]$ is the maximized log-likelihood of sites in $\boldsymbol{y}$ and with the maximum likelihood estimates (MLE) of parameters on that same data and $lnL[\boldsymbol{z}, \hat{\theta}(\boldsymbol{y})]$ is the predicted log-likelihood of sites in $\boldsymbol{z}$ with parameters set to the MLEs based on sites $\boldsymbol{y}$ ($lnL\left[\boldsymbol{z}, \hat{\theta}(\boldsymbol{z})\right] and\ lnL[\boldsymbol{y}, \hat{\theta}(\boldsymbol{z})]$ are defined similarly).

To test model fit, half of the alignment sites were selected at random as a training set while the remaining were used as the test set, resulting in non-overlapping sets of equal sizes. This was done four more times, resulting in five total partitions. The roles of the training and testing sets were then switched to calculate CV2. We estimated the CV2 value for all partitions with all combinations of models (LG, C60, UDM64, MEOW(60,20), MEOW(40,40), MEOW(80,00)) and trees (all MLE trees estimated from the full dataset by each model). Table 1 summarizes the best CV2 values for each partition and model over all trees. In all cases, the MEOW(60,20) model appears to fit the best of all tested models for each partition and so it was used for all subsequent analyses.

*Table 1. The resulting 2-fold Cross-Validation for the five different partitions and the calculated mean. For each model, tree, and partition, CV2 values were estimated, but here we report only the best CV2 value per partition and model. The best value per column is shown in bold in the table.*

| Model | Partition 1 | Partition 2 | Partition 3 | Partition 4 | Partition 5 | Mean CV2 |
|---|---|---|---|---|---|---|
| LG | -1,763,095 | -1,763,145 | -1,763,095 | -1,763,144 | -1,763,128 | -1,763,121 |
| C60 | -1,681,909 | -1,681,891 | -1,681,896 | -1,681,900 | -1,681,900 | -1,681,899 |
| UDM64 | -1,676,842 | -1,676,854 | -1,676,866 | -1,676,873 | -1,676,880 | -1,676,863 |
| MEOW(60,20) | **-1,666,284** | **-1,666,557** | **-1,666,333** | **-1,666,413** | **-1,666,673** | **-1,666,452** |
| MEOW(40,40) | -1,669,007 | -1,668,866 | -1,668,933 | -1,669,000 | -1,668,944 | -1,668,950 |
| MEOW(80,00) | -1,670,380 | -1,670,268 | -1,670,182 | -1,670,531 | -1,670,545 | -1,670,381 |

**Partitioning dataset based on compositional heterogeneity.** First, the counts of all 20 amino acids were tabulated across an alignment for all taxa, including missing data when a taxon is absent. A chi-squared test of the null hypothesis that composition is homogeneous over taxa [4] was performed on these counts using the chisq.test() function in R.The Pearson's residuals for each taxon and amino acid from the chi-squared test were extracted from the output of the chisq.test() object in R. In this case, for each amino acid (aa) and taxon, the Pearson's residual for that amino acid is calculated as:

$$R_p = \frac{O - E}{\sqrt{E}}$$

Where O is the observed count of the amino acid for that taxon and E is the expected count assuming homogeneous amino acid usage over taxa. If a taxon lacked a protein, each amino acid residual for that taxon was computed as missing data. These per-taxon residuals were collated together for all alignments into a 93-row, 20,000-column matrix where each row represented an alignment in the dataset, and each column represented an amino acid in a taxon (20 amino acids * 100 taxa = 20,000 columns). We treated this matrix of residuals as independent data distributions which can be clustered using a Gaussian mixture model [5]. Ten independent clusterings were performed using the "gaussian_pk_sj" model in the R package MixAll, which allowed for free weighting of cluster proportions and free variance among variables but equal variance within clusters (i.e., a diagonal covariance matrix). The number of clusters k was set to 5, to ensure artefacts arising from partitioning were kept to a minimum [6,7]. The criterion for model selection was set to BIC and a SemiSEM clustering strategy was used to handle missing data. The highest-scoring clustering run according to BIC across the ten independent clustering runs was chosen as the partitioning strategy to be used for this dataset.

The chosen partitioning strategy produced 5 partitions of roughly equal gene and site totals, where partitions 3 and 4 were predominantly composed of nuclear-encoded/mitochondrial-localized genes and/or mitochondrial-encoded genes, and the other partitions were predominantly composed of nuclear-encoded genes (Table 2).

*Table 2. Gene content for each of the generated partitions. Genes are organized by the compartment in which they are encoded, and the total number of sites and average length of the alignment excluding gaps for each partition are reported.*

| **Partition** | **Nuclear** | **Mitochondrial + Nuclear** | **Mitochondrial** | **Total genes** | **Total sites** | **Avg. Length Non-Gaps** |
|---|---|---|---|---|---|---|
| **1** | 119, 1974, 21231, 2942, 3008, 3252, 49839, 90211, atpC, clpP, Der1-2, sucA | atp9, rpl20 | atp4, nad1, rps8 | **19** | **4597** | **3330** |
| **2** | 1092, 1378, 2282, 2811, 418, 4684, 5434, 5518, 8768, 9, Der11, Der3, Der5, Der6, Der7 | atp3, rpl27, sdh2 | atp8 | **19** | **4993** | **3726** |
| **3** | 1140, 15631, 15911, 27742 | atp1, nad10, nad7, nad8, rpl11, rpl1, rpl2, rps14, rps2, tufA | cob, rpl31 | **16** | **3212** | **2547** |
| **4** | 1052, hslV | cox11, nad11, nad2, nad4l, nad9, rpl16, rpl6, rps19, rps4 | atp6, ccmFC, cox1, cox3, nad3, nad4, nad5, nad6, tatC | **20** | **4661** | **3383** |
| **5** | 15632, 16172, 2000, 21468, 22332, 3060, 342, 3735, 4140, 72, 778, 8455, 8754, Der13, ksgA, lpd, ndufaf5, trmE | cox2 | N/A | **19** | **4999** | **3388** |
| **Totals** | **53** | **25** | **15** | **93** | **22462** | **16376** |

**Selecting optimum amino acid compositions to model for each partition.** The binomial test of two proportions gives initial sets, $\mathcal{G}$ and $\mathcal{F}$, of amino acids that are enriched or depleted in the two groups of taxa. These sets were further refined as follows. From these initial sets, we constructed subsets, $\mathcal{G}'$ and $\mathcal{F}'$ of $\mathcal{G}$ and $\mathcal{F}$; we considered all possible subsets excluding the empty sets. For each subset, $\mathcal{G}'$ and $\mathcal{F}'$and each of the two groups of taxa, we obtained counts of the total number of $\mathcal{G}'$amino acids and $\mathcal{F}'$amino acids in the partition. This resulted in a $2\ x\ 2$ table of counts that was used to get a chi-square test statistic for the test of the null hypothesis that the probability of an amino acid being in $\mathcal{G}'$, given that it is either in $\mathcal{G}'$ or $\mathcal{F}'$, is homogeneous over the two groups.

When this test statistic is large it suggests that heterogeneity of amino acid usage of amino acids in $\mathcal{G}'$ and $\mathcal{F}'$. Excluding amino acids not in these two groups makes it easier to detect the important subsets. Note that the degrees of freedom for the test are constant over choices of subsets $\mathcal{G}'$ or $\mathcal{F}'$, regardless of the sizes of each subset. The chi-squared test statistic was retained for each combination. Combinations were ranked by their chi-squared test statistic and the top 50 combinations were retained for GFmix analysis.
